# A Unified Framework for TCR-pMHC Structural Model Assessment

**DOI:** 10.1101/2025.10.09.681411

**Authors:** Alex Ascunce-París, Roc Farriol-Duran, Miguel Romero-Durana, Alfonso Valencia, Víctor Guallar

**Affiliations:** Barcelona Supercomputing Center (BSC), Barcelona, Spain; Catalan Institution for Research and Advanced Studies (ICREA), Pg. Lluís Companys 23, 08010 Barcelona, Spain

## Abstract

T cell receptor (TCR) recognition of peptide-MHC complexes (pMHCs) is a central determinant of cellular immunity. Structural insights are key to understanding TCR-pMHC interactions, however progress is limited by the scarcity of experimental structures (PDB = 232 TCR-pMHC class I complexes, date 01/2025). Although protein modelling tools have rapidly advanced, their accuracy on TCR-pMHC complexes is hampered by CDR loop hypervariability, conformational flexibility, and the lack of reliable quality assessment without experimental references. To evaluate the performance of recent protein modelling algorithms on TCR-pMHC complexes, we benchmarked three general-purpose (AlphaFold3, Boltz-2, Chai-1) and two TCR-specific (tFold-TCR, TCRmodel2) modelling tools using a benchmark set of 20 experimentally determined PDB structures, demonstrating AlphaFold3 superior performance. To expand structural coverage beyond the existing experimental landscape, we present a framework that enables quality assessment of TCR-pMHC structural models without requiring experimental reference structures. To this end, we integrate multiple modelling and interface confidence metrics (pLDDT, ipTM, iPAE, pDockQ) into explainable random forest classifiers trained on 1160 models of 232 experimentally determined PDB TCR-pMHC class I complexes. This approach reliably distinguishes low-, acceptable-, medium-, and high-quality models, outperforming the combination of these metrics using literature-established thresholds. Leveraging our quality tier framework, we used AlphaFold3 to generate and evaluate the largest synthetic dataset of TCR-pMHC class I structural models to date from VDJdb-annotated sequences, comprising 33,820 complexes (169,100 models, 5 models per complex) and representing a >70-fold expansion of available structures. We demonstrate the applicability of our quality tier stratification in three settings: 1) filtering validating vs. non-validating TCR-pMHC interaction in sequence databases (VDJdb), 2) enrichment of biologically validated interactions versus synthetic negatives amongst higher quality complexes, and 3) enhancing the predictive performance of TCR-pMHC pairing models. Altogether our TCR-pMHC structural quality tier framework provides a scalable and interpretable approach for improving modelling and functional analyses of TCR-pMHC class I complexes, with translational applications in T-cell and TCR-based immunotherapies.

## Introduction

Recognition of peptide-MHC complexes (pMHCs) by T cell receptors (TCRs) is a central event in cellular immunity [1]. Most TCRs are αβ heterodimers that mediate antigen recognition through six hypervariable loops, known as complementarity-determining regions (CDRs). Each chain contains three CDRs, which together simultaneously engage both the presented peptide or epitope and the MHC molecule [2]. To maximize coverage of potential epitopes, human TCR repertoires are generated by V(D)J recombination, a somatic gene rearrangement process that combines variable (V), diversity (D), and joining (J) gene segments with random nucleotide insertions and deletions at the junctions. This process is estimated to produce over 10^11^ distinct receptors in an individual, out of approximately 10^15^ potential possibilities, prior to thymic selection, providing each individual with extraordinary clonal and structural diversity in T cell responses [3].

Structural insights into TCRs and their cognate pMHC partners are essential for understanding the geometry and biophysics of the TCR-pMHC interface, which is governed by recognition specificity, and ultimately drives T-cell activation [4–6]. Although recent deep-learning and AI-based methods have attempted to predict TCR-pMHC interactions from sequence data, these approaches often fail to generalize to “unseen” epitopes, those not represented in any of the TCR-pMHC complexes within the training sets [7]. Incorporating biophysical information derived from structural data could capture binding motifs governing molecular recognition [8,9], ultimately improving the generalizability of predictive models to novel epitopes [6,10,11]. However, the experimental determination of TCR-pMHC structures remains challenging, with only a few hundred TCR-pMHC class I complexes deposited in the PDB [12] (n=232, date 01/2025) .

Recent advances in protein structure prediction have opened new opportunities for modelling TCR-pMHC complexes using existing sequence datasets with annotated validations. Protein modelling tools provide predictions of molecular complexes with varying accuracy, especially for immune proteins such as antibodies and TCRs [13,14]. However, the intrinsic sequence variability and conformational flexibility of CDR loops, combined with the limited number of experimentally determined TCR-pMHC structures and the absence of co-evolutionary constraints, pose major challenges for these algorithms [6,13]. Examples include general-purpose tools such as AlphaFold3 [15], Chai-1 [16], Boltz-2 [17], and TCR-specific TCRmodel2 [13] and tFold-TCR [18].

Evaluating the quality of these predictions in the absence of experimentally determined structures remains difficult due to the lack of reliable quality metrics that do not rely on comparison with reference experimental structures. Several confidence scores such as pLDDT, ipTM, iPAE [19], and pDockQ (v1 and v2) [20,21] have been proposed, yet their success remains limited in some biological contexts, particularly immune complexes [22]. Beyond these modeling challenges, the annotation quality of the TCR-pMHC complexes in sequence databases can further limit predictive accuracy. Sequence databases such as VDJdb [23] have been reported to contain a substantial fraction of false positives or interactions of uncertain confidence, particularly among immunodominant epitopes [24]. This issue can compromise the predictive performance of both sequence-based and structure-based TCR-pMHC pairing predictors trained on this data. [24,25].

In this study, we benchmarked state-of-the-art general and TCR-specific protein modeling tools on a representative set of TCR-pMHC class I complexes from the PDB [12]. Based on these results, we developed an explainable, reference-free framework to assess structural model quality using random forest classifiers trained on AlphaFold3 [15] models derived from all available PDB [12]-annotated TCR-pMHC class I experimental structures. The models were grouped into four quality tiers based on comparisons with their corresponding experimental reference structures, and these quality tiers were then predicted using model confidence metrics as features. We applied this framework to generate a large in-house dataset of 33,820 AlphaFold3 [15] TCR-pMHC class I models from VDJdb [23]-annotated sequences. Finally, we demonstrate that the resulting quality tiers can be used to (i) filter out non-validating entries from immunodominant epitopes in sequence databases such as VDJdb, (ii) enrich biologically validated interactions within higher-quality tiers, and (iii) enhance the predictive performance of TCR-pMHC pairing models.

## Results

### 1. AlphaFold3 provides reliable structural models for TCR-pMHC class I complexes with experimentally resolved PDB structures

We first evaluated the performance of different state-of-the-art (SOTA) modelling tools: AlphaFold3 [15], Chai-1 [16], Boltz-2 [17], TCRmodel2 [13] and tFold-TCR [18] on a benchmark set of 20 experimentally resolved TCR-pMHC class I structures obtained from the PDB [12]. The benchmark set included 10 complexes deposited before (pre) and 10 after (post) the training cutoff dates for the five modelling tools.

Model quality was assessed by comparing the predicted structural models to their corresponding experimental structures using the Cα interface RMSD of the TCR chains (TCR-iRMSD) and the DockQ score [26], categorizing models into quality tiers as follows: high quality (HQ) if TCR iRMSD < 2 Å and DockQ > 0.8; medium quality (MQ) if TCR iRMSD < 5 Å and DockQ between 0.49 and 0.8; acceptable quality (AQ) if TCR iRMSD < 5 Å and DockQ between 0.23 and 0.49; and low quality (LQ) if TCR iRMSD > 5 Å and DockQ < 0.23 (Fig 1A). We adopted this four-tier classification implementing the thresholds defined by the DockQ metric [26] for bona-fide protein-protein interactions. For each complex, AlphaFold3 [15], TCRmodel2 [13] and Chai-1 [16], generated five models, while Boltz-2 [17] and tFold-TCR [18] generated one model per complex. For the former tools, the best model for each complex was selected based on the closest match to the corresponding experimental structure.

**Fig. 1.**
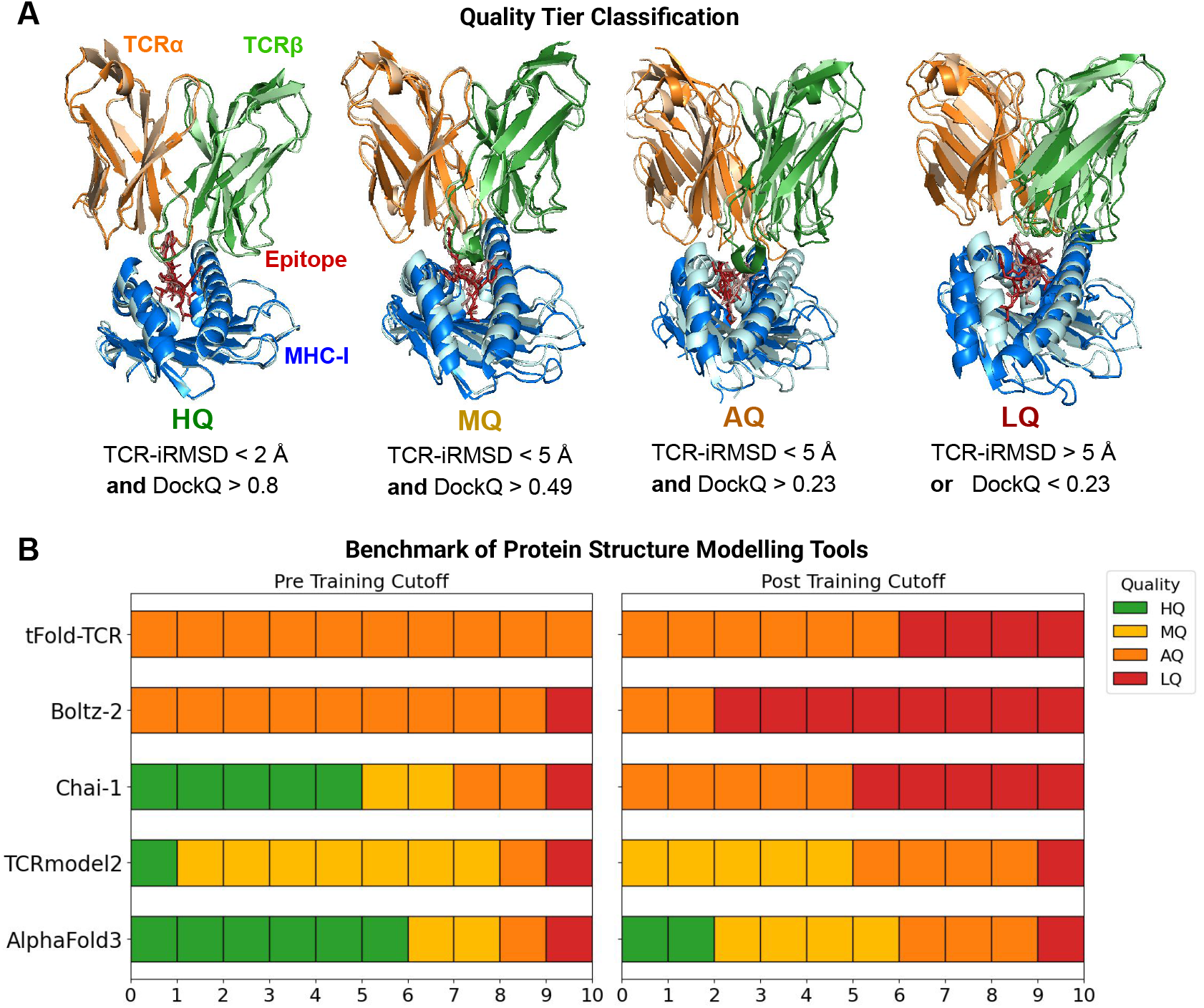
Benchmarking of TCR–pMHC Structure Prediction Models. (A) Criteria used to classify model quality into tiers: HQ (high quality), MQ (medium quality), AQ (acceptable quality), and LQ (low quality), based on TCR interface RMSD (TCR-iRMSD) and DockQ score thresholds. The structures display superimpositions of the crystal structure (dark colors) versus the corresponding AlphaFold3 model (light colors). **(B)** Benchmarking of state-of-the-art structure prediction tools; general-purpose (AlphaFold3, Chai-1, and Boltz2) and TCR-specific (TCRmodel2 and tFold-TCR) on 20 TCR-pMHC crystal structures. This dataset was divided between 10 crystals deposited at the PDB before (pre) and 10 after (post) the training cutoff dates of the tools.

AlphaFold3 outperformed the other modelling tools, with 90% of evaluated complexes above the acceptability threshold across both pre- and post-training test sets (Fig. 1B). TCRmodel2 achieved comparable acceptability rates but with overall lower qualities: only 1 HQ pre- and no HQ post-cutoff models, compared with 6 HQ pre- and 2 HQ post-cutoff models for AlphaFold3. Chai-1 showed similar performance to AlphaFold3 in the pre-training test set, with one fewer HQ model and one more AQ model, but exhibited low performance in the post-training set, with 50% of complexes classified as LQ and the other 50% as AQ. Boltz-2 and tFold-TCR underperformed in both test sets, reaching only AQ levels without any MQ or HQ models. In the pre-training set, 9 models of Boltz-2 were AQ, compared with 10 for tFold-TCR, while in the post-training set Boltz-2 reached only 2 AQ models, and tFold-TCR had 6. Although Boltz-2 and tFold-TCR achieved low iRMSDs in the pre-training set (<2 Å) and reasonable iRMSDs in the post-training set (<5 Å), their DockQ scores remained low (Supplementary Fig. S1), placing their models in lower-quality tiers and resulting in overall underperformance compared with the other three tools.

Taken together, these results suggest that AlphaFold3 provides accurate structural predictions for TCR-pMHC class I complexes within this benchmark set of 20 representative structures, with 90% of models classified as acceptable, including 20% high-quality (HQ), 40% medium-quality (MQ), and 30% acceptable-quality (AQ) models. This represents a clear improvement over TCRmodel2, the next best-performing tool, which yielded 0% HQ, 50% MQ, 40% MQ models.

To further evaluate model quality, we generated AlphaFold3 [15] structural models for all or all TCR-pMHC class I structures currently available in the PDB [12]. We next compared them to their corresponding experimental structures. The dataset was again stratified based on deposition date relative to the AlphaFold3 training cutoff (09/2021), resulting in 173 complexes (865 models) deposited prior to the cutoff (pre) and 59 complexes (295 models) deposited afterwards (post).

Full interface, chain-specific interface and CDRα/β RMSDs (iRMSD, TCR-iRMSD, peptide-RMSD, MHC-iRMSD, CDRsα-RMSD, CDRsβ-RMSD) revealed that a substantially larger fraction of the models in the pre test set achieved <2 Å compared to post models. Nevertheless, in the post-test set most RMSDs remained <5 Å, indicating that predictions still captured overall geometry with reasonable accuracy (Fig 2A, left and center panel). A similar trend was observed for DockQ scores: pre models were skewed toward higher values, while post models shifted slightly lower, though the majority remained above the 0.23 acceptability threshold (Fig. 2A, right panel).

**Fig. 2.**
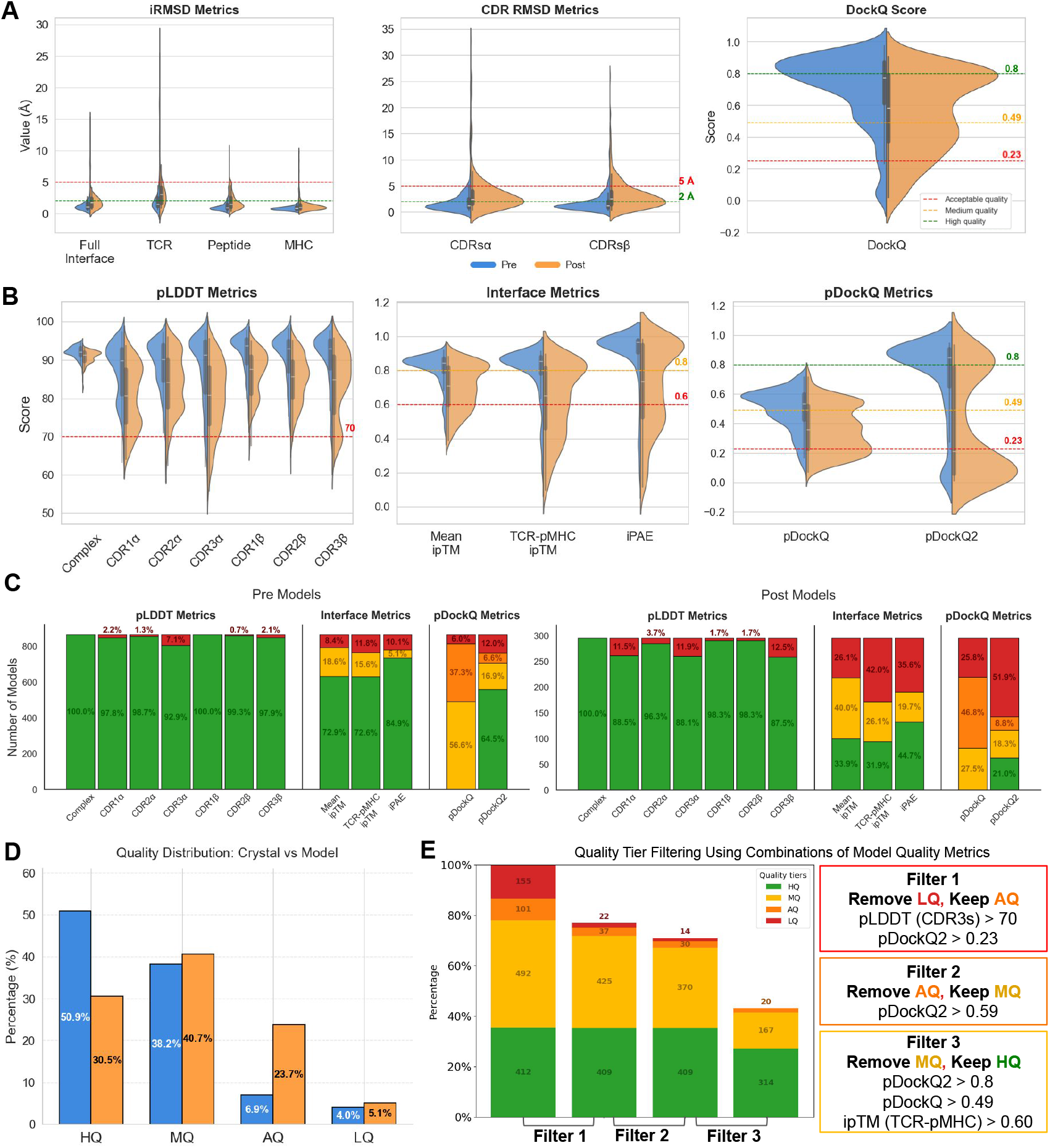
Evaluation of TCR–pMHC Class I Structural Models. (A) Distribution of crystal vs. model comparison metrics of structural models generated from sequences of experimentally solved PDB TCR-pMHC crystals, divided into pre- and post-test sets according to whether they were deposited before or after the AlphaFold3 training cutoff date (09/21). Metrics include interface RMSD for the full interface, TCR chains, peptide and MHC chains, mean RMSD of the three CDRs from both α and β chains, and the DockQ score. Red lines indicate acceptability thresholds (2 Å RMSD and 0.23 DockQ); yellow lines denote medium-quality thresholds (0.49 DockQ); green lines indicate high-quality thresholds (2 Å RMSD and 0.8 DockQ). **(B)** Distribution of model quality metrics for the pre- and post-test sets: pLDDTs of the full complex and the six individual CDRs, interface metrics (mean ipTM, TCR–pMHC ipTM, and iPAE), and predicted DockQ scores (pDockQ and pDockQ2). Red lines indicate acceptability thresholds (70 pLDDT, 0.23 pDockQ, and 0.6 for interface metrics); yellow lines denote medium-quality thresholds (0.49 pDockQ and 0.8 for interface metrics); green lines represent high-quality thresholds (0.8 pDockQ). **(C)** Effect of applying default model-quality metric thresholds. Models were classified into HQ (green), MQ (yellow), AQ (orange), or LQ (red) according to the defined criteria. **(D)** Distribution of model quality in the pre- and post-test sets according to crystal vs. model comparison metrics, using the thresholds defined in Figure 1A. **(E)** Effect on model-quality distribution after applying different filters of combinations of model-quality metrics to the dataset.

Based on these comparisons, models were assigned to previously defined quality tiers (Fig. 1A). Within the pre-cutoff set, 27% were HQ, 21% MQ, 13% AQ, and 39% LQ. In the post-cutoff set, 9% were HQ, 44% MQ, 5% AQ, and 41% LQ (Fig. 2D). These results indicate that, although overall accuracy decreases in the post-cutoff test set, most notably with a shift from HQ to MQ, a substantial fraction of models still reach acceptable or better quality highlighting the utility of AlphaFold3 for generating reliable TCR-pMHC models from sequence data.

### 2. Evaluation of protein modelling and interface confidence metrics for classifying TCR-pMHC class I structural model quality

To enable large-scale generation of synthetic structural data from annotated TCR-pMHC complexes, we aimed to develop a reference-free quality tier classification system based on model confidence metrics. To this end, we evaluated multiple metrics that can be directly extracted from structural predictions without requiring corresponding experimental structures. These metrics were considered at multiple levels: full-complex structural accuracy (pLDDT) and a subset focusing exclusively on CDRα/β-specific pLDDTs; standard protein-protein interface confidence metrics such as mean ipTM as well as TCR-pMHC-specific ipTM and average iPAE and predicted DockQ scores (v1 and v2) [20,21] at the TCR-pMHC interface.

As shown in Fig. 2B, left panel, full-complex pLDDTs obtained on most of TCR-pMHC structural models were consistently high (means ∼92% pre, ∼90% post). CDR3 pLDDTs <70% were uncommon, though post-test distributions shifted lower by ∼5-10% on average. ipTM and iPAE values were generally reduced in the post-test set, with TCR-pMHC-ipTMs close to the 0.6 acceptability threshold (Fig 2B, center panel). Predicted DockQ (pDockQ) systematically underestimated true DockQ scores obtained from the corresponding experimental structure comparison, and both pDockQ and pDockQ2 displayed bimodal distributions, particularly in the post-test set, reflecting a mixture of higher- and lower-quality modelled structures (Fig. 2B, right panel).

We then applied literature-established thresholds for each metric to classify structural model quality. These default thresholds revealed substantial discrepancies in quality classification across different metrics within both the pre- and post-test datasets (Fig. 2C). Accounting for global and CDR-level pLDDTs, nearly all structural models in the pre-test set were HQ (>95%), with minimal LQ fractions (<5%), whereas in the post-test set, HQ proportions remained dominant (≈85–95%) but with a modest increase in LQ counts. Interface-based metrics were more variable: ipTM and TCR-pMHC ipTM ranked ≈70% in the pre-test set as HQ, dropping to ≈30% HQ in the post-test set, with corresponding increases in AQ and LQ fractions. iPAE followed a similar but slightly more optimistic trend (≈85% HQ pre, ≈45% HQ post). In contrast, pDockQ classified almost no HQ models in either set (≈6% pre, ≈25% post), while pDockQ2 yielded the largest LQ fractions overall (≈12% pre and >50% post). These results demonstrate that applying default thresholds produces highly inconsistent quality distributions across metrics.

We next evaluated whether combining these metrics leveraging established thresholds could better discriminate between quality tiers while prioritizing the most accurate TCR-pMHC structural models amongst AlphaFold3 [15] predictions. Using exhaustive metric combinations in a 5-fold cross-validation scheme, we derived integrative filters with very high discrimination capacity (Table 1, Fig. 2D). Specifically, distinguishing LQ from AQ (Filter 1; composed of pLDDT CDR3s > 70 and pDockQ2 > 0.23) achieved a precision of 90%, while distinguishing AQ from MQ (Filter 2; pDockQ2 > 0.59) reached a precision of 94%, albeit with lower recall (87% for both). In contrast, discriminating HQ from MQ (Filter 3; pDockQ2 > 0.8, pDockQ > 0.49, and ipTM TCR-pMHC > 0.60) was less robust, yielding an HQ-enriched but not pure subset (68% precision, 37% recall).

**Table 1.** Model Quality Metric-based filters applied to classify model quality tiers. Filters and corresponding metric threshold combinations used to remove models belonging to specific quality tiers (LQ, AQ, MQ), as illustrated in Figure 2E. Each filter represents the set of model-quality thresholds applied to exclude models of the target tier. The associated precision and recall values indicate the performance of each metric combination in correctly identifying models in PDB-modeled structures.

| Filter | Remove | Keep | Best Metric Combination | Precision | Recall |
| --- | --- | --- | --- | --- | --- |
| 1 | LQ | AQ | pLDDT (CDR3s) > 70<br>pDockQ2 > 0.23 | 0.98 | 0.87 |
| 2 | AQ | MQ | pDockQ2 > 0.49 | 0.95 | 0.87 |
| 3 | MQ | HQ | pDockQ2 > 0.8<br>pDockQ > 0.49<br>ipTM (TCR-pMHC) > 0.60 | 0.69 | 0.35 |

When applying these filters sequentially to the full dataset (Fig. 2E), the first filter (Filter 1) removed nearly all LQ models (∼86%) but also eliminated ∼60% of AQ models and ∼14% of MQ models, reflecting the high precision but low recall values. The second filter (Filter 2) had minimal additional effect because most AQ models were already removed (reducing AQ from 37 to 30) but also removed ∼13% of remaining MQ models. Finally, the third filter (Filter 3) removed ∼60% of remaining MQ models and ∼25% of HQ models, achieving only partial enrichment for HQ and highlighting the stringent nature of sequential filtering steps.

Together, these findings underscore the limitations of relying on individual metrics, despite being combined in optimized scoring schemes, to capture the diverse properties of modeled TCR-pMHC structures. Such an approach requires the use of heterogeneous thresholds and is prone to systematic biases. These results highlight the need for a unified, reference-free framework capable of accurately and consistently classifying TCR-pMHC model quality, thereby enabling the generation of large-scale synthetic TCR-pMHC complex datasets.

### 3. Random forest classifiers integrating protein modelling and interface confidence metrics robustly distinguish structural quality tiers of TCR-pMHC structural models

To address these challenges, we developed a unified, reference-free method to accurately assess TCR-pMHC structural model quality. We trained three independent random forest classifiers to distinguish between adjacent quality tiers: LQ vs. AQ, AQ vs. MQ, and MQ vs. HQ. Using separate binary classifiers simplifies the learning task, allowing each model to focus on the subtle differences between two neighboring tiers. This design also enhances interpretability, as each classifier emphasizes the most informative metrics for distinguishing one tier from the next.

Training was performed using quality labels derived from structural model vs corresponding PDB-experimental structures comparisons as ground truth (Fig. 1A), and model confidence metrics as features. By using the same set of metrics previously described (complex-pLDDT and CDRα/β-specific pLDDTs mean ipTM, TCR-pMHC-specific ipTM, average iPAE and predicted DockQ scores v1 and v2 of the TCR-pMHC interface), the classifiers leverage both global and interface-specific information to discriminate between adjacent quality tiers.

When distinguishing LQ from AQ (RF LQ-AQ) and AQ from MQ (RF AQ-MQ), the random forest classifiers achieved precision comparable to that obtained using default threshold combinations (96% for both classifiers vs. 98% and 95% with default thresholds), while recall was substantially improved (97% and 98% vs. 87% for both with default thresholds). For the MQ from HQ separation (RF MQ-HQ), both precision and recall were lower than for the other tiers but still surpassed the performance of default thresholds (precision 73% and recall 75% vs. 69% and 35%, respectively) (Table 1 vs. Table 2).

**Table 2.**
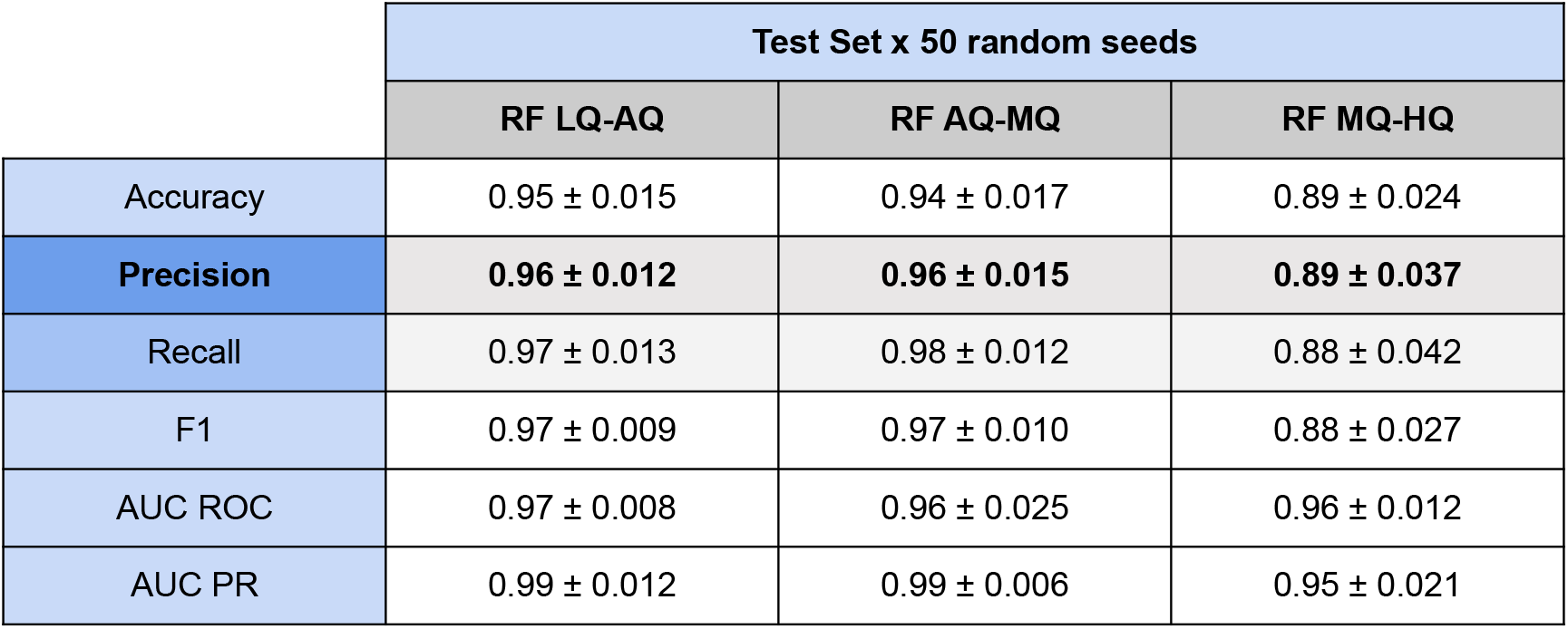
Performance metrics of Random Forest classifiers across consecutive quality tiers. Performance metrics for the three Random Forest classifiers trained using models classified by consecutive quality tiers: RF LQ-AQ, RF AQ-MQ, and RF MQ-HQ. Reported metrics include Accuracy, Precision, Recall, F1-score, Area Under the ROC Curve (AUC ROC), and Area Under the Precision-Recall Curve (AUC PR). Metrics were evaluated on the test set using 50 random seeds, and values are reported as mean ± standard deviation.

|  | Test Set x 50 random seeds |  |  |
| --- | --- | --- | --- |
|  | RF LQ-AQ | RF AQ-MQ | RF MQ-HQ |
| Accuracy | 0.95 ± 0.015 | 0.94 ± 0.017 | 0.89 ± 0.024 |
| **Precision** | **0.96 ± 0.012** | **0.96 ± 0.015** | **0.89 ± 0.037** |
| Recall | 0.97 ± 0.013 | 0.98 ± 0.012 | 0.88 ± 0.042 |
| F1 | 0.97 ± 0.009 | 0.97 ± 0.010 | 0.88 ± 0.027 |
| AUC ROC | 0.97 ± 0.008 | 0.96 ± 0.025 | 0.96 ± 0.012 |
| AUC PR | 0.99 ± 0.012 | 0.99 ± 0.006 | 0.95 ± 0.021 |

To further understand feature contributions, we utilized SHAP (SHapley Additive exPlanations) values to interpret each model’s predictions. SHAP values provided a clear indication of which features played the most significant roles in driving the model’s predictions [27] (Fig 3A). Across all tasks, pLDDT scores were the most informative features. For LQ-AQ, α-chain CDRs, especially CDR3α, contributed most, followed by β-chain CDRs and the global complex pLDDT. pDockQ2 and iPAE provided moderate contributions, while ipTM had lower but notable impact. For AQ-MQ, CDR1α and CDR2α gained importance relative to CDR3α, and β-chain CDRs showed a more balanced profile. ipTM of the TCR-pMHC interface became more relevant, with pDockQ and iPAE remaining secondary yet significant. In MQ-HQ, α- and β-chain CDR pLDDTs contributed nearly equally and strongly, with complex pLDDT, pDockQ, and iPAE supporting predictions; ipTM remained less influential but consistently represented.

**Fig. 3.**
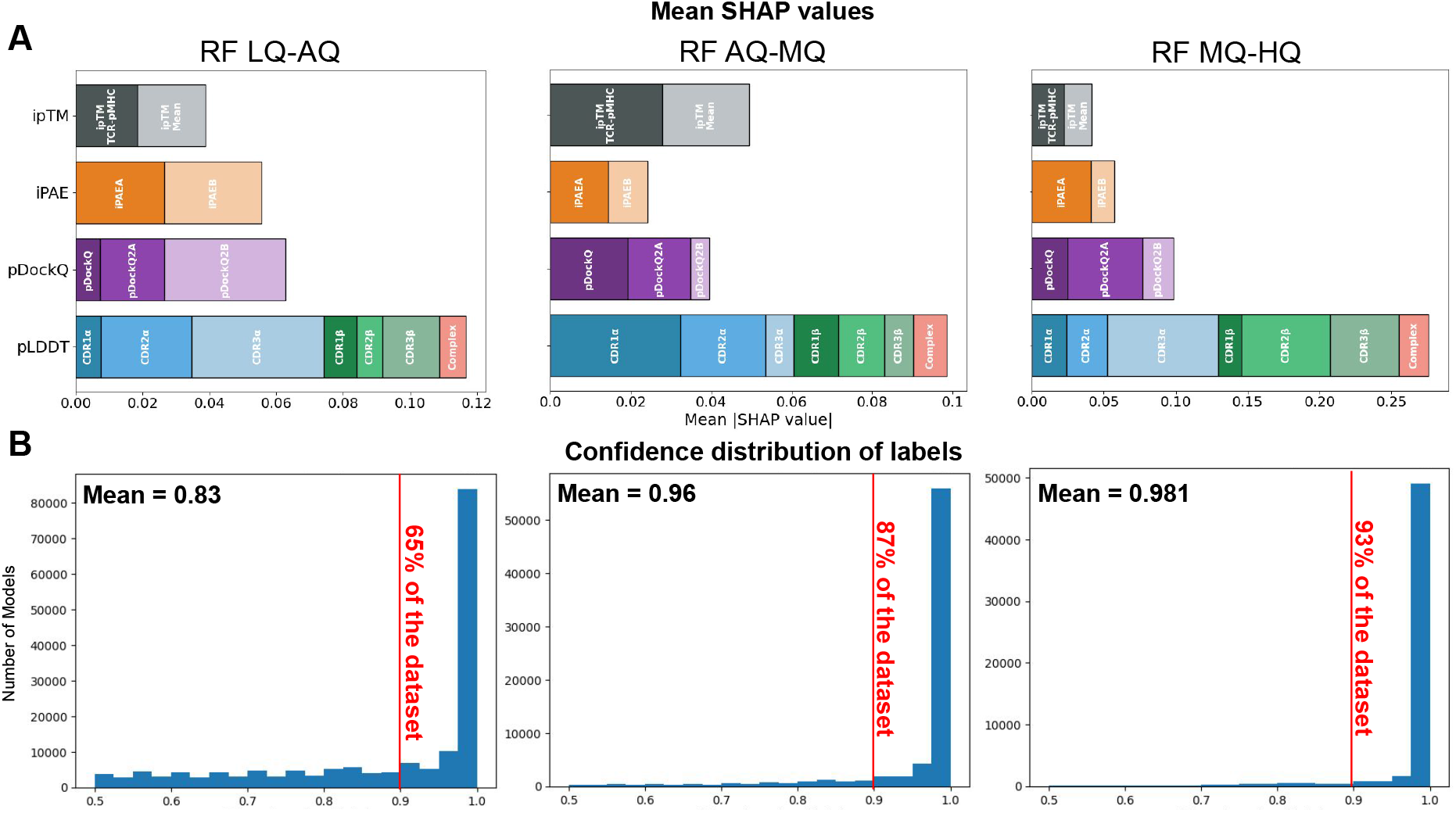
Explainability analysis of Random Forest Classifiers. (A) Shapley feature importance for model-quality metrics, including pLDDT (full complex, CDRs of the α chain, and CDRs of the β chain), ipT (mean and TCR-pMHC interface), iPAE, pDockQ, and pDockQ2. This analysis highlights which features contribute most to the Random Forest predictions of structural quality tiers. (B) Histograms showing the confidence distribution of the quality label predictions for each model, illustrating how confidently each Random Forest classifier assigns structural quality tiers across the synthetic dataset.

Finally, we assessed the confidence of the assigned labels via bootstrap analysis. The resulting confidence scores indicated high reliability for all labels. When classifying AQ-LQ, the mean confidence was 0.83, with 65% of labels above 0.9 (Fig. 3B). Confidence was even higher for MQ-AQ, with a mean of 0.96 and 87% of labels exceeding 0.9, and for MQ-HQ, with a mean of 0.98 and 93% of labels above 0.9. Although the HQ vs. MQ classifier showed lower precision, it was highly confident in its predictions.

In summary, our results demonstrate that a reference-free, multi-feature approach can reliably distinguish TCR-pMHC class I structural model quality tiers. pLDDT scores within the CDRs consistently provide the strongest predictive signal, while additional metrics such as pDockQ, iPAE, and ipTM enhance model discrimination, particularly for more challenging transitions. The classifiers also produce highly confident label assignments, supporting their potential utility in guiding downstream structural analyses and model selection.

### 4. Expanding the TCR-pMHC class I structural landscape through AlphaFold3-based modelling and quality stratification of the entire VDJdb

To expand the structural landscape of TCR-pMHC class I complexes, we generated an in-house large-scale synthetic structural dataset. Leveraging AlphaFold3 [15] we modelled sequences annotated in the VDJdb [23] resulting in 33,820 unique TCR-pMHC class I complexes, with five models per complex, yielding a total of 169,100 modelled structures.

Once modelled, we can observe in Fig. 4 that AlphaFold3 structures from our synthetic models from VDJdb data generally exhibit lower quality scores compared to those predicted from sequences with experimental structures already available at the PDB, including those in our post-training cutoff test set (Fig. 2B). Although the post-PDB structures were not part of AlphaFold3’s training set and thus would represent unseen data, their predicted qualities are substantially higher. In the synthetic structural dataset, pLDDTs generally remain above 70 and are comparable to the AlphaFold3 post-test set (∼90 for the full complex and ∼80 for the CDRs) (Fig. 4A, left panel). However, interface metrics (Fig. 4A, center panel), including both mean and TCR-pMHC-specific ipTMs and iPAE, are consistently skewed toward lower values, with means near the acceptability threshold of 0.6. Some complexes show particularly high iPAE values exceeding 0.8, although few exhibit similarly high ipTM values. Finally, pDockQ, and especially pDockQ2 (Fig. 4A, right panel), show very low values, with only a few complexes reaching medium-quality thresholds for pDockQ and acceptable-quality thresholds for pDockQ2.

**Fig. 4.**
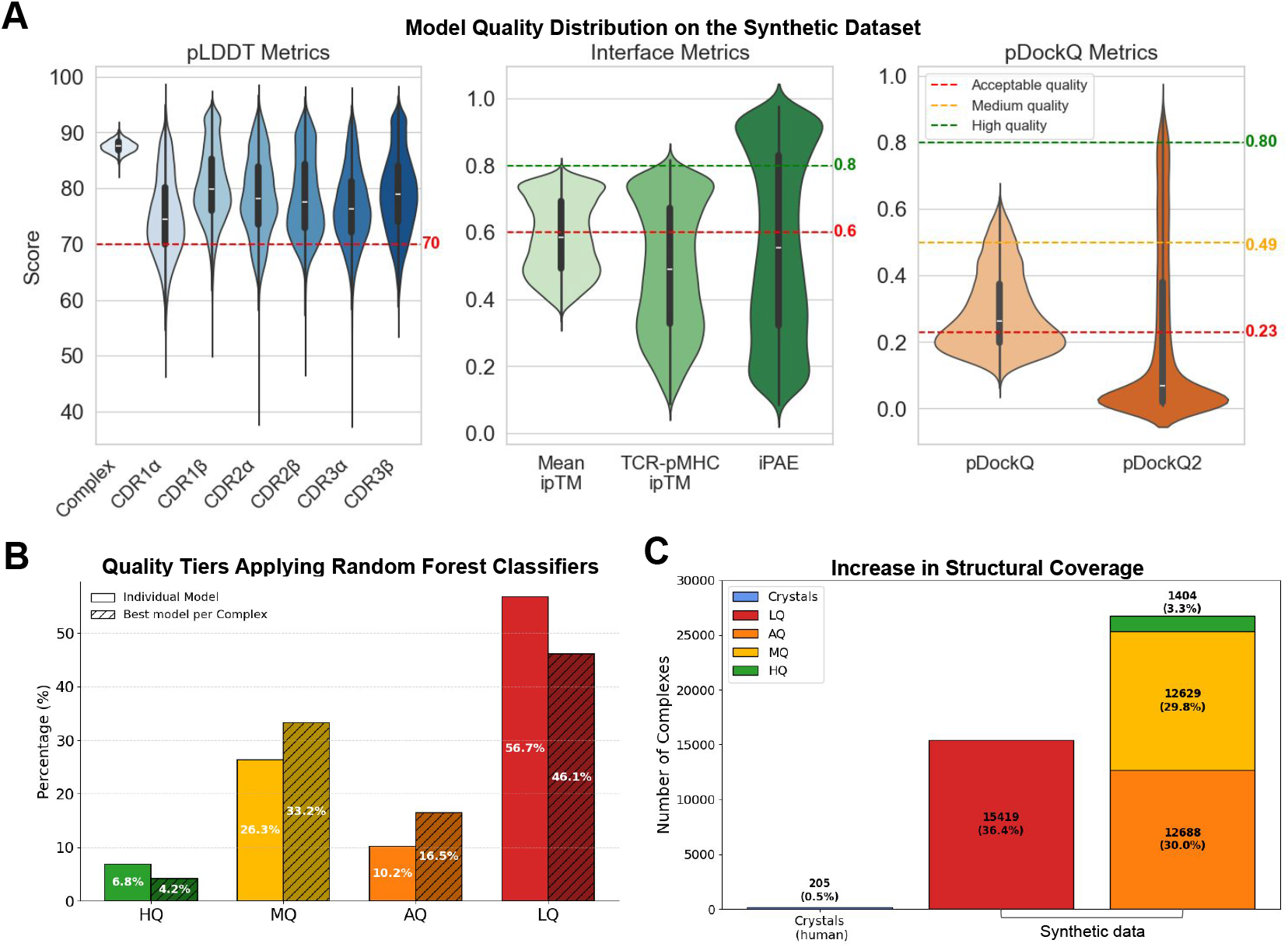
Expanding the TCR-pMHC class I Structural Landscape. (A) Distribution of model quality metrics for the synthetic dataset: pLDDTs for the full complex and the six individual CDRs; interface metrics including mean ipTM, TCR–pMHC ipTM, and iPAE; and predicted DockQ scores (pDockQ and pDockQ2). Red lines indicate acceptability thresholds (70 pLDDT, 0.23 pDockQ, 0.6 for interface metrics), yellow lines indicate medium-quality thresholds (0.49 pDockQ, 0.8 for interface metrics), and green lines indicate high-quality thresholds (0.8 pDockQ). **(B)** Quality distribution of the synthetic dataset generated with AlphaFold3: percentages are shown per individual model (smooth bars) and per complex, aggregating the scores from the five models generated per input query (hatched bars). **(C)** Quality classification in the final structural dataset combining AlphaFold3 models and PDB crystal structures.

To achieve a consistent quality classification of the obtained modelled-structures, we applied our random forest-based framework, assigning all synthetic TCR-pMHC structural models into four quality tiers. Models were categorized in the following percentages: HQ (6.8%), MQ (26.3%), AQ (10.2%), and LQ (56.7%) (Fig. 4C). Dissecting the results and analyzing the best model per complex decreased the proportion of low-quality predictions (HQ 4.2%, MQ 33.2%, AQ 16.5%, and LQ 46.1%), reflecting variability across AlphaFold3 solutions (Fig. 4C, hatched bars). Notably, compared to modelled PDB experimental structures in the post-test set (Fig. 2D), again, the synthetic dataset exhibits substantial differences in quality distribution, with a marked increase in LQ models (from 5.1% in the post test set to 56.7%) and corresponding decreases in HQ, MQ, and AQ models (30.5%, 40.7%, and 23.7% in the post test set vs. 6.8%, 26.3%, and 10.2%).

The lower scores and overall reduced quality classification in our synthetic dataset relative to the post-training cutoff test set suggest that experimentally resolved structures represent a restricted and relatively homogeneous subset of the TCR-pMHC structural landscape, whereas the synthetic models capture a broader and more diverse structural space. Although nearly half of the generated models were classified as low structural quality, this approach enabled a substantial expansion of the TCR-pMHC available structural landscape: from 232 initial experimental structures (205 human complexes) deposited in the PDB [12] to 18,401 acceptable- to high-quality synthetic models, representing a >70-fold increase i n the structural space for TCR-pMHC class I complexes (Fig. 4D). This expanded dataset enhances the structural representation of the TCR-pMHC annotated triad sequences, providing a reliable resource for downstream applications, including predictive modelling, epitope mapping, and structural analyses.

### 5. Quality tier stratification effectively identifies non-validating TCR-pMHCs in VDJdb

To investigate whether modelled-structural quality alone can be used to filter TCR-pMHC complexes within the VDJdb [23] recently reevaluated as non-validating, we leveraged the dataset provided by Messemaker et al., TCRvdb [24]. This dataset was generated testing the two of the most studied epitopes at VDJdb [23] GLCTLVAML (GLC, Epstein-Barr virus, EBV) and YLQPRTFLL (YLQ, SARS-CoV-2) paired with multiple TCRs and of the HLA-A*02:01-restricted in an harmonized T-cell validation system. This allowed the identification of TCR-pMHC complexes with experimentally validating and non-validating interactions. From our synthetic dataset, we extracted the structural models corresponding to these TCR-pMHC pairs and assessed whether validating interactions (padj < 1 × 10^−5^) were enriched among higher-quality models (Fig. 5A).

**Fig. 5.**
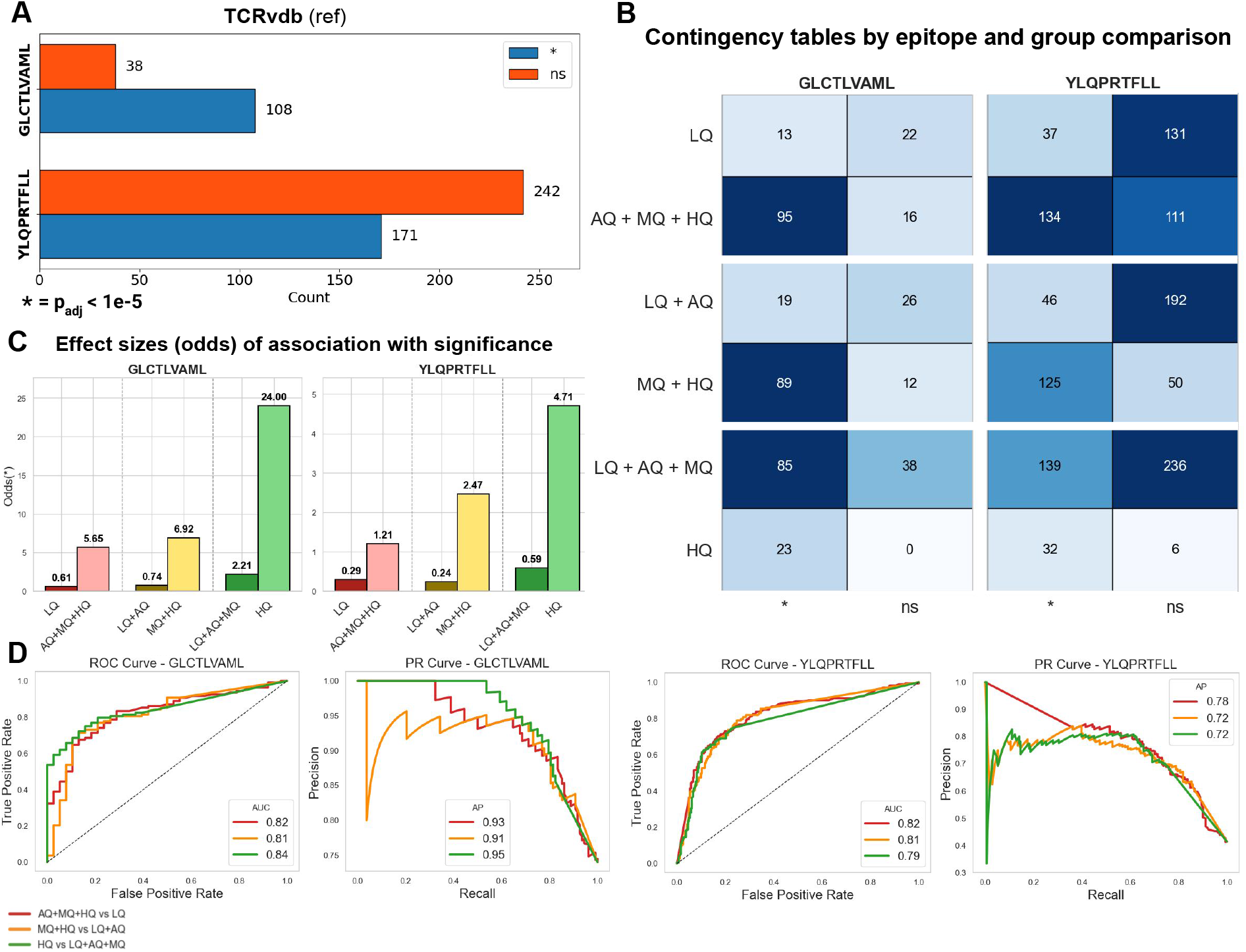
Quality Tier Stratification Identifies False Positives in VDJdb. (A) TCRvdb counts of significant () versus non-significant (ns) instances for the epitopes GLCTLVAML and YLQPRTFLL (padj < 1×10^−5^). **(B)** Contingency tables show quality-based splits of the dataset and their significance ( / ns) for both epitopes. **(C)** Effect sizes (odds ratios) indicate the association between each quality combination and significance (*) for both epitopes. **(D)** AUC-ROC and AUC-PR summarize the ability of quality tiers to distinguish significant from non-significant instances for both epitopes.

We first assessed the relationship between model quality labels (LQ, AQ, MQ, HQ) and the significance of the interaction using Fisher’s exact tests and bonferroni correction for multiple comparisons (padj < 0.017). For the GLCTLVAML epitope, acceptable-quality models (AQ+MQ+HQ) were significantly enriched for validation TCR-pMHCs compared to low-quality models (LQ) (odds ratio = 5.65 CI 95% (3.37-9.46) vs 0.61 CI 95% (0.311.18), p < 0.0001). Restricting to medium- and high-quality models (MQ+HQ) further increased enrichment (odds ratio = 6.92 CI 95% (3.87-12.38) vs 0.74 CI 95% (0.42-1.32) for LQ+AQ, p < 0.0001), while high-quality models alone (HQ) exhibited the strongest association with validating interactions (odds ratio = 24.0 CI 95% (3.25-177.41), p = 0.003). Notably, in this last scenario, lower-quality models (LQ+AQ+MQ) still retained a moderate association with significance (odds ratio = 2.21 CI 95% (1.51-3.22)), indicating that a substantial number of positive instances are not captured by the HQ label (Fig. 5B and 5C).

A similar trend was observed for the YLQPRTFLL epitope: acceptable-quality models (AQ+MQ+HQ) were enriched for validating TCR-pMHCs (odds ratio = 1.21 CI 95% (0.94-1.55) vs 0.29 CI 95% (0.20-0.41) for LQ, p < 0.0001), medium- and high-quality models (MQ+HQ) showed increased enrichment (odds ratio = 2.47 CI 95% (1.78-3.42) vs 0.24 CI 95% (0.18-0.33) for LQ+AQ, p < 0.0001), and high-quality models alone (HQ) achieved again the highest enrichment (odds ratio = 4.71 CI 95% (2.09-10.66) vs 0.59 CI 95% (0.48-0.73), p < 0.0001).

In parallel, we evaluated the discriminative power of our random forest predicted probabilities using ROC and PR curves. For GLCTLVAML, ROC AUC values ranged from 0.81–0.84 and PR AUC values from 0.91–0.95 across scenarios, with no statistically significant differences, reflecting robust discrimination between validating and non-validating interactions (Dolong’s test + Bonferroni correction, padj < 0.017). AUC PR at low recall was particularly high for higher-quality models, indicating effective prioritization of true positives and alignment with the contingency table enrichment.

For YLQPRTFLL, ROC AUC ranged from 0.79–0.82 and PR AUC from 0.72–0.78. Discrimination was generally consistent across model qualities, though acceptable-quality models (AQ+MQ+HQ) exhibited higher PR AUCs compared with LQ models (Bootstrap padj < 0.05).

Together, these analyses demonstrate two complementary perspectives. On the one hand, enrichment analysis shows that higher-quality models are more likely to capture validating TCR-pMHCs amongst the two most studied epitopes at VDJdb. Medium- and high-quality models (MQ+HQ) offer a balanced strategy, minimizing false positives while retaining most validating TCRpMHCs interactions, whereas HQ models maximize the likelihood of true positives but reduce recall. On the other hand, AUC ROC and AUC PR metrics confirm that the associated probabilities provide reliable discrimination between validating and non-validating TCRpMHCs reevaluated by Messemaker et al., [24]. These results reinforce the utility of our model quality framework both as a filtering criterion and a quantitative predictor of TCRpMHC interactions.

### 6. High-quality structural models enrich for positive TCR-pMHC complexes in the IMMREP23 dataset and e enhance predictive performance of TCR-pMHC pairing methods

We next investigated whether modelled-structure quality influences the predictive capacity of TCR-pMHC pairing algorithms. We used the IMMREP23 TCR-pMHC dataset from Kaggle [28], which comprises 3,483 TCR-pMHC class I complexes representing 20 epitopes with an approximate 1:5 positive-to-negative ratio. Negative examples were generated by pairing peptides with TCRs binding to unrelated epitopes [29].

We successfully modelled 3,480 TCR-pMHC complexes from this dataset using AlphaFold3 [15], extracted model quality metrics, and stratified structures into quality tiers using our random forest classifiers. This resulted in 596 positive and 2,884 negative TCR-pMHCs distributed unevenly across the quality tiers (Fig. 6A). We observed that as model quality increased, both the number and proportion of positive instances increased (from 9.9% in the LQ tier to 65.6% in the HQ tier), whereas negative instances decreased (from 90.1% to 34.4%), yielding a positive-to-total ratio increase from 0.10 in AQ to 0.66 in HQ (Fig 6C, left).

**Fig. 6.**
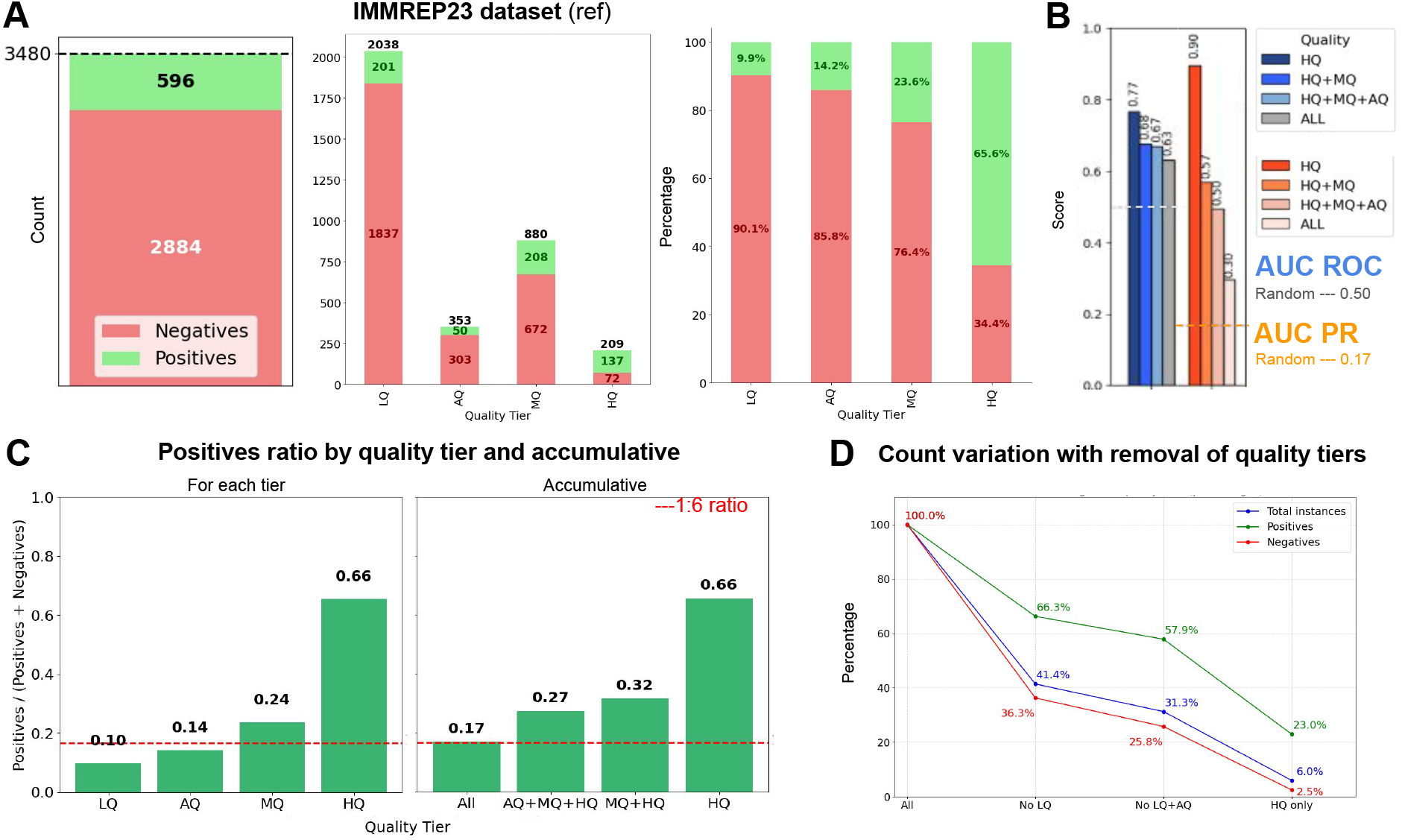
High-Quality Structural Models Improve TCR-pMHC Pairing. (A) Overview of the IMMREP23 dataset, showing the ratio of positive versus negative instances in the full dataset, stratified by quality tier and expressed as percentages. **(B)** AUC-ROC and AUC-PR of TCoaRse, our in-house structure-based TCR-pMHC pairing method (unpublished), for pairing prediction across different quality tiers in the IMMREP23 dataset; dashed lines indicate random performance. **(C)** Increase in the positive-to-total ratio for each quality tier and cumulatively when progressively removing lower-quality structural models in the IMMREP23 dataset. **(D)** Changes in the percentages of total instances (blue), positives (green), and negatives (red) when progressively removing lower-quality tiers in the IMMREP23 dataset.

Considering the entire dataset and progressively removing lower-quality tiers, the positive-to-negative ratio increased from the initial 1:6 (0.17) to 4:6 (0.66), indicating that higher-quality structures are enriched in paired TCR-pMHCs, thus biologically plausible (Fig. 6C, right). Notably, removing LQ structures significantly reduced the number of instances to test (from 3480 to 1441), while retaining 66% of positives (394 positives from 596 total). Removing AQ models further decreased total instances by ∼10% (1089 instances) while reducing positives by only 8% (345 instances). Excluding MQ structures reduced the total dataset from 100% to 6% (209 instances), with positives decreasing to 23% (137 instances) (Fig. 6D). The HQ tier is predominantly populated by positive instances, whereas MQ strikes the optimal compromise between precision and recall, consistent with the results of Section 5.

To evaluate the performance of our structure-based predictor, TCoaRse (an in-house tool, unpublished; manuscript in preparation), we tested it on this dataset stratified by structural quality tiers. We observed that predictive performance improved as the quality of the structures increased. Specifically, the AUC ROC increased from 0.63 across the full dataset to 0.67, 0.68, and 0.77 when progressively excluding the lowest-quality structures. Similarly, the AUC-PR increased from 0.30 to 0.50, 0.57, and 0.90. These improvements were statistically significant according to the DeLong test with multiple Bonferroni correction (padj < 0.05).

These results indicate that high-quality structural models enhance predictive performance and that quality tier stratification is a powerful approach to enrich for biologically meaningful TCR-pMHC interactions while reducing false positives.

## Discussion

This study introduces a reference-free framework to assess the quality of TCR-pMHC class I modeled structures, addressing a key gap in the field: evaluating the quality of structural predictions in the absence of experimentally determined structures. Previous approaches have largely focused on creating or benchmarking individual structural quality metrics, both for overall protein prediction accuracy (pLDDT) and for interface-specific metrics (ipTM, iPAE, or pDockQ). However, these metrics were developed in broader protein structure prediction contexts and are not specifically tailored to TCR-pMHC complexes. Consequently, their performance and interpretability can vary depending on the structural or biological context in which they are applied, particularly in immune receptor complexes, where the high flexibility of the CDR loops poses significant challenges for accurate quality estimation [13,22]. In contrast, our framework integrates multiple structural quality metrics into a simple quality tier system obtained using explainable machine learning classifiers, providing more reliable and standardized assessments of the quality of TCR-pMHC modelled structures.

In addition, we introduce the largest synthetic TCR-pMHC class I structural dataset to date, generated using AlphaFold-multimer version 3.0.0 [15] to model complexes annotated in VDJdb [23]. This dataset represents a major resource for immunology research, vastly expanding the structural coverage of the TCR sequence space by more than 70-fold compared to existing structures at the PDB [12]. Future efforts will focus on making this dataset publicly available through a dedicated database to facilitate access and further research. We also plan to expand the dataset, increasing its coverage and incorporating TCR-pMHC class II complexes.

The combination of this large-scale dataset and our modeled structure quality classification pipeline enables the identification of immunologically meaningful TCR-pMHC interactions and provides a practical mechanism to filter TCR-pMHC sequence databases such as VDJdb [23] retaining only high-confidence pairs, such as experimentally validating TCR-pMHC triads [24]. Moreover, by enriching medium- and high-quality models, our approach enhances the accuracy of downstream TCR-pMHC pairing predicting frameworks, offering tangible benefits for applications such as TCR-based cancer immunotherapies [30].

Furthermore, our results show that model quality metrics are consistently lower for modeled VDJdb [23] structures than for modeled experimentally solved structures deposited in the PDB [12], even when these structures were not included in AlphaFold3’s training set (Fig. 4A). This reflects the fact that TCR-pMHC class I structures in the PDB [12] are drawn from a relatively narrow region of sequence space, where experimentally resolved complexes tend to resemble each other in both sequence and structure. In contrast, the broader and lower-scoring distributions observed in the synthetic dataset underscore the limited experimental coverage and highlight the critical importance of generating large-scale, diverse structural models. Such diversity is essential to explore novel TCR-pMHC combinations and expand our understanding of the full immunological repertoire, which is not captured by the currently resolved, highly homogeneous structures.

Despite the advances encompassed by our work, important limitations remain. High-confidence predictions are constrained by the intrinsic flexibility of CDR loops, which are difficult to model accurately due to AlphaFold3’s rigid-body treatment of complexes [31] that does not fully capture the dynamic conformational states critical for recognition. These challenges are further compounded by the limited availability of experimental crystal structures. Future work could integrate molecular dynamics simulations or multiple AlphaFold3 modelled structures starting from multiple seeds per complex to better capture conformational ensembles, as recent studies suggest that AlphaFold3 models can capture alternative binding conformations [32]. Consequently, although nearly half of the synthetic structural models were classified as low quality, they may still provide valuable structural insights for downstream applications, as they can represent binding modes that exist but are not yet captured in current databases and are therefore considered low-confidence.

Overall, our framework demonstrates that combining structural modeling with data-driven quality assessment enhances both the structural and functional interpretation of TCR-pMHC interactions. Beyond improving the selection of modeled structures, it provides a scalable approach for enriching TCR-pMHC structural datasets, reducing non-validated yet annotated-as-positive TCR-pMHC entries in VDJdb [23], and guiding immunoinformatic predictors for TCR pairing. Altogether, our framework and dataset offer tangible benefits for both basic research and translational applications.

## Methods

### 1. Data retrieval

Sequence data was retrieved from VDJdb database [23] (downloaded 17/01/2025). We restricted our dataset to human MHC class I TCR-pMHC complexes with four-digit resolution allele annotation, paired CDR3α and CDR3β, and fully annotated V and J genes (TRAV, TRAJ, TRBV, and TRBJ). Complete TCR information was required to reconstruct the variable domain sequence using Stitchr [33]. These entries were then submitted to ANARCI [34] for annotation of individual CDR regions using the IMGT numbering scheme. CDR1 was defined as positions 27–38, CDR2 as positions 56–65, and CDR3 as positions 105–117 in the alignment. Duplicated TCRα /TCRβ combinations were removed, and only CDR3 sequences starting with a C and ending with a phenylalanine F were retained according to IMGT definition of CDR3 loops [35].

Experimentally resolved TCR-pMHC class I crystal structures were retrieved from the TCR3D database [36,37], a curated collection of TCR-related structures derived from the Protein Data Bank (PDB) [12] downloaded 08/01/2025).

The TCRvdb dataset was obtained from Messemaker et al. [23]. From this dataset, we retained 614 instances with annotated padj values, of which 559 were also present in our VDJdb data. Following the authors’ recommendation, a threshold of 1×10^−5^ was used to define statistical significance.

Finally, the IMMREP23 dataset [28,29] was downloaded from the official GitHub repository:https://github.com/justin-barton/IMMREP23/.

### 2. Benchmarking of protein modelling tools on TCR-pMHC class I complexes

We benchmarked state-of-the-art protein modelling tools: AlphaFold3 [15], and Chai-1 [16], Boltz-2 [17], TCRmodel2 [13] and tFold-TCR [18]. A benchmark set of 20 TCR-pMHC class I complexes with known crystal structures was curated from the PDB [12]. These complexes were selected to include an equal number of structures deposited before (Pre PDB IDs 1oga, 1qsf, 2bnq, 1qrn, 2vlr, 1ao7, 2ckb, 1mi5, 1bd2, 4jry) and after (Post PDB IDs 8qfy, 8en8, 8wul, 8wte, 8f5a, 8i5c, 8enh, 8eo8, 8shi, 8gom) the training cutoff dates of the modelling tools to allow for an unbiased evaluation of their generalization capabilities.

Each tool was used to predict the structures of the selected complexes. Predicted models were then compared to the corresponding crystal structures using multiple metrics: Cα interface root mean square deviation (iRMSD) of all chains of the complex, heavy atoms RMSD of the CDRs, DockQ score [26], and TCR crossing and incident angles [36]. The interface was defined as all atoms within 10 Å of the peptide chain.

Models were classified based on two representative metrics: TCR chain iRMSD (TCR iRMSD) and DockQ score as follows: high quality (HQ) if TCR iRMSD < 2 Å and DockQ > 0.8; medium quality (MQ) if TCR iRMSD < 5 Å and DockQ between 0.49 and 0.8; acceptable quality (AQ) if TCR iRMSD < 5 Å and DockQ between 0.23 and 0.49; and low quality (LQ) if TCR iRMSD > 5 Å and DockQ < 0.23 (Fig 1A).

### 3. Structural modelling with AlphaFold3

We generated TCR-pMHC class I structural models using AlphaFold3 [15] from sequences of experimentally determined PDB TCR-pMHC class I complexes as well as from sequences previously retrieved and processed from the VDJdb database [23] and from the IMMREP23 dataset (ref, ref). Our synthetic dataset comprises 33,820 TCR-pMHC complexes, with 5 models per complex (169100 models). Modelling was performed using AlphaFold3 version 3.0.0 on a setup with 1 GPU and 20 CPUs, with an average computation time of approximately 1.5 hours per complex.

### 4. Model quality metrics

Model quality was evaluated using a set of complementary metrics spanning multiple structural levels. Full-complex structural accuracy was assessed using predicted local distance difference test (pLDDT) scores, while CDR-specific accuracy was evaluated by computing pLDDTs for individual CDRα and CDRβ loops to capture local modeling quality. Protein-protein interface metrics included the mean predicted interface TM-score (ipTM), reflecting global interface confidence. For the TCR-pMHC interface specifically, we computed interface-specific ipTM, average interface predicted aligned error (iPAE) [13,19], and predicted DockQ scores (v1 and v2) [20,21] to evaluate interface geometry and accuracy. These metrics were used both individually and in combination to assess structural quality and to train the Random Forest classifiers for assigning models to discrete quality tiers.

### 5. Structural annotation and model processing

Chains of the PDB structures were annotated using the script **“mir-1.0-SNAPSHOT.jar”** provided by Karnaukov et al. [10] which outputs a general.txt file containing the correspondence between chain type and chain ID for each PDB complex. Model chains were automatically annotated during modeling with the following chain assignments: A: MHC-I, B: β2-microglobulin, C: peptide, D: TCRα, E: TCRβ. Resulting CIF files were subsequently converted to PDB format using BeEM [38]. To calculate TCR-pMHC-specific DockQ, average iPAE, pDockQ, and pDockQ2 scores, chains were merged such that chain A included MHC-I, β2-microglobulin, and the peptide, while chain B included TCRα and TCRβ. The pDockQ [20] and pDockQ2 [21] scripts were obtained from their original publications and adapted for use with AlphaFold3 output.

### 6. Random Forest Classifiers

Classifiers were trained on 232 PDB [12] complexes modelled with AlphaFold3 [15] with 5 models per complex (1,160 models in total). Quality labels were derived from crystal vs. corresponding model comparison metrics (TCR-iRMSD and DockQ, See Methods Section 2, Fig 1A) applied to experimentally resolved PDB structures. Input features consisted of crystal-independent model quality metrics, including full-complex pLDDTs, CDRsα/β-specific pLDDTs, mean predicted interface TM-scores (ipTM) and TCR-pMHC ipTM, average interface predicted aligned error (iPAE) [19] and and predicted DockQ scores (v1 and v2) [20,21] for the TCR-pMHC interface.

Three independent Random Forest classifiers were trained to distinguish between adjacent quality tiers: LQ vs. AQ, AQ vs. MQ, and MQ vs. HQ. Training was performed using scikit-learn with default parameters and evaluated through a stratified 80/20 train-test split repeated over 50 random seeds to ensure robustness and stability of predictions. Data splits were designed so that no models from the same complex appeared in both training and test sets, preventing data leakage. Feature importance was quantified using SHAP values [27], and the most informative metrics were subsequently employed to select the best model per complex for downstream analyses. To evaluate the reliability of the assigned quality labels, a bootstrap-based confidence estimation approach was applied.

### 7. Enrichment analysis

Enrichment of biologically validated interactions within structural model quality tiers was assessed using Fisher’s exact test for contingency tables and AUC ROC and AUC PR for continuous scores. Since our Random Forest classifiers were trained to discriminate adjacent quality tiers (LQ vs AQ, AQ vs MQ, MQ vs HQ), we calculated chained conditional probabilities to assign each model a probability for all four quality tiers. Probabilities were computed sequentially as follows: P(LQ) and P(AQ) were derived directly from the LQ vs AQ classifier.

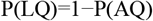

For models that have a probability of being MQ, P(AQ) was adjusted to reflect that some AQ-classified models are actually MQ:

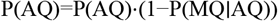

P(MQ) was calculated as the probability of being MQ given that the model was previously classified as AQ.

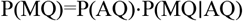

For models that have a probability of being HQ, P(MQ) was adjusted to reflect that some MQ-classified models are actually HQ:

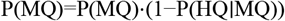

P(HQ) was calculated as the probability of being HQ given that the model was previously classified as MQ

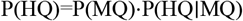

Finally, probabilities were normalized so that:

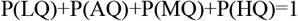

This approach ensures continuous and interpretable probability estimates for all four tiers, avoids missing values (NaNs), and allows computation of ROC and PR curves for each quality category comparison and statistical significance.

### 8. Statistical analysis

Receiver Operating Characteristic (ROC) curves and the corresponding Area Under the Curve (AUC ROC) values were evaluated using the Dlong test to compare predictive performance across models.

Precision-Recall (PR) curves and the associated Area Under the Curve (AUC PR) values were assessed using bootstrap resampling. Bonferroni correction was applied to account for multiple comparisons in both cases. All statistical tests were two-sided, and significance thresholds were adjusted accordingly, with an adjusted p-value < 0.05 considered statistically significant.

## Data availability

The scripts used to evaluate the models can be found at https://github.com/Alexasparis/AF3TCRpMHC.git

## GRAPHICAL ABSTRACT

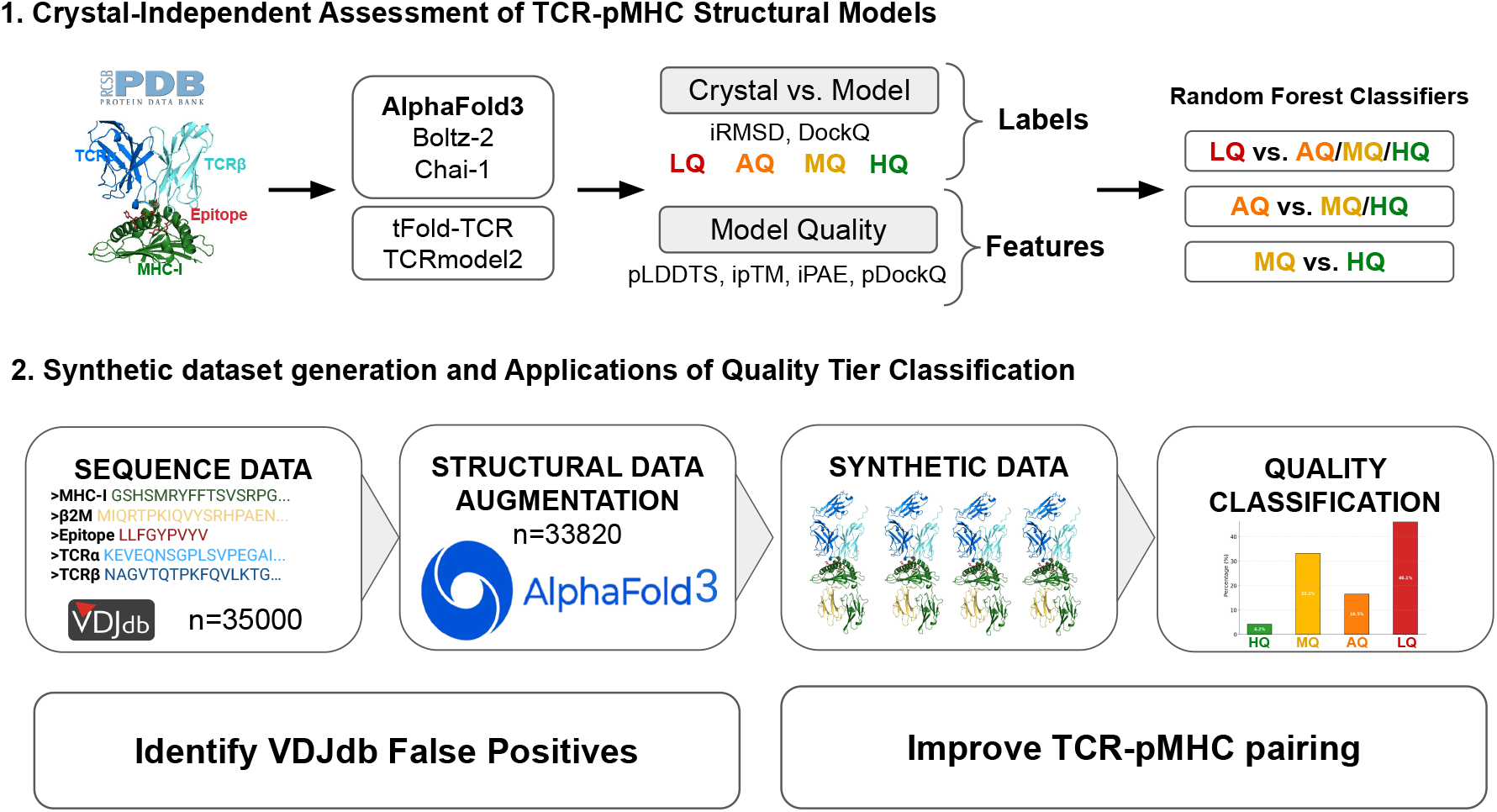

**Fig. S1.**
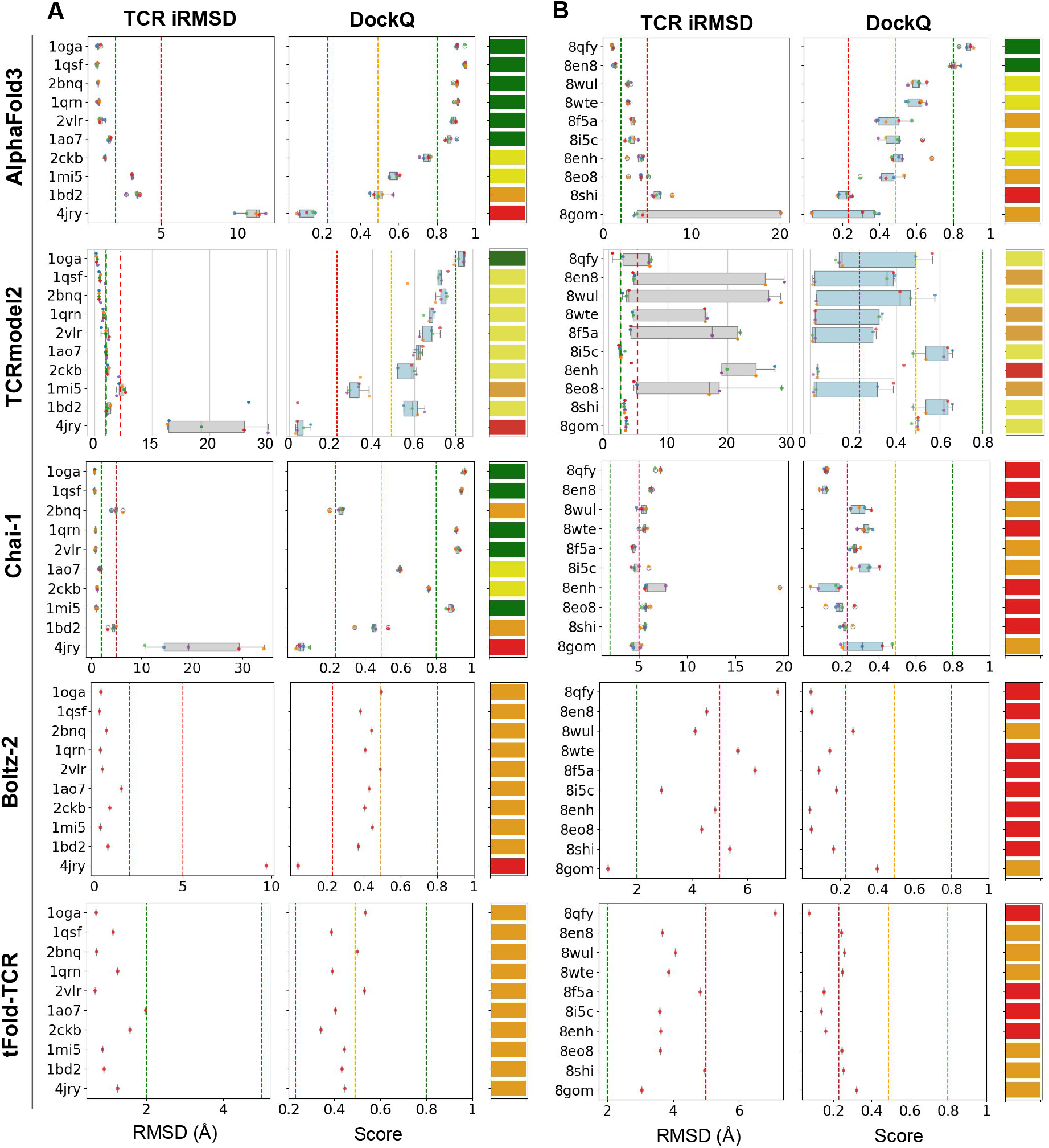
Benchmarking of AlphaFold3, Chai-1, Boltz-2, TCRmodel2, and tFold-TCR. The figure shows interface Cα iRMSD of the TCR chains, DockQ scores, and quality tier classifications for TCR-pMHC class I structural models. Quality thresholds for model classification were defined in Figure 1A, and models were assigned to tiers based on these thresholds. (A) iRMSD and DockQ values for 10 crystal structures deposited in the PDB before the training cutoff dates of the modelling tools. (B) iRMSD and DockQ values for 10 crystal structures deposited after the training cutoff.

